# Patterning ganglionic eminences in developing human brain organoids using morphogen gradient inducing device

**DOI:** 10.1101/2023.06.20.545705

**Authors:** Narciso Pavon, Karmen Diep, Feiyu Yang, Rebecca Sebastian, Beatriz Martinez-Martin, Ravi Ranjan, Yubing Sun, ChangHui Pak

## Abstract

In early neurodevelopment, the central nervous system is established through the coordination of various neural organizers directing tissue patterning and cell differentiation. Better recapitulation of morphogen gradient production and signaling will be crucial for establishing improved developmental models of the brain *in vitro*. Here, we developed a method by assembling polydimethylsiloxane (PDMS) devices capable of generating a sustained chemical gradient to produce patterned brain organoids, which we termed <u>M</u>orphogen-gradient <u>I</u>nduced <u>B</u>rain <u>O</u>rganoids (MIBOs). At 3.5 weeks, MIBOs replicated Dorsal-Ventral patterning observed in the Ganglionic Eminence (GE). Analysis of matured MIBOs through single-cell RNA sequencing revealed distinct Dorsal GE derived CALB2+ interneurons (INs), Medium Spiny Neurons (MSNs), and MGE derived cell types. Finally, we demonstrate long term culturing capabilities with MIBOs maintaining stable neural activity in cultures grown up to 5.5 months. MIBOs demonstrate a versatile approach for generating spatially patterned brain organoids for embryonic development and disease modeling.

**Motivation:** The use of small molecules for the guided differentiation of brain organoids has proven to be a useful tool for modeling various aspects of early neurodevelopment. Still, the embryonic brain is patterned through a combination of morphogen gradients stemming from organizer regions. To address this inconsistency, we designed a device capable of mimicking neural organizers by maintaining a steady morphogen gradient. We used this device to expose forebrain organoids to multiple gradient conditions and cataloged the diversity of cell types produced.

## Introduction

Early human neurodevelopment relies on a wide variety of signaling factors to accurately and efficiently pattern the brain. Prior studies in model systems revealed that the diffusion and reaction-diffusion of various morphogens can instruct neural induction, and subsequently, cell fate patterning, leading to the establishment of the Dorsal-Ventral (D-V) and Anterior-Posterior (A-P) axes of the central nervous system (CNS). At the most anterior region of the developing neural tube, the primary brain vesicle prosencephalon further develops into telencephalon which encompasses the cerebrum ^1, 2^. The telencephalon itself produces a wide range of diverse cell types in a D-V dependent manner ^3^. The dorsal most regions of the telencephalon become the cortex and the ventral regions give rise to the subpallium.

Transiently, during the embryonic stages of neurodevelopment, the subpallium holds the ganglionic eminences (GEs) comprising the medial GE (MGE), lateral GE (LGE), and caudal GE (CGE). The GEs contain ventral ventricular zones (VZs) responsible for the production of GABAergic Interneurons (INs) and projection neurons ^4–6^. The GABAergic cell types go on to innervate a wide variety of regions in the telencephalon depending on place of origin. The MGE produces INs that will migrate into the cortex and striatum ^6–11^. The LGE generates Medium Spiny Neurons (MSNs) for the striatum and INs that will migrate into the olfactory bulb ^12–16^. Finally, the CGE produces INs that will travel to the striatum, cortex, and amygdala ^13, 17, 18^. While much of what we know of the GEs has come from animal models, recent characterizations of post-mortem human fetal brain tissue have aided in enlightening the unique transcriptomes of the human GEs ^19–23^. Due to the unique progenitor pools of the GEs, reproducing the anatomical regionalization in GEs is a key component of accurately modeling brain development.

To facilitate further investigation into the complexity of human neurodevelopment, in recent years, 3D human pluripotent stem cell (hPSC)-derived cell cultures, termed neural organoids, have been developed to serve as models for various regions of the CNS depending on the protocol used ^24–38^. Particularly, forebrain organoids have been well established using guided differentiation techniques to generate region-specific organoids modeling single brain regions such as the cortex, striatum, MGE, and optic cup, respectively ^27, 28, 39–42^. To model the connectivity of multiple embryonic brain regions, organoids representing different brain regions or other tissues have been derived separately, before fusing to form assembloids ^39–41, 43–45^. While proven useful, current protocols do not fully recapitulate the early neural patterning processes.

At present, protocols that enable the generation of D-V or A-P axes in a single organoid through establishment of morphogenic gradients are extremely limited. To overcome these limitations, novel methods have been developed including the chimeric organoid model which fused an aggregate of Sonic hedgehog (Shh) expressing cells to a forebrain organoid, thus mimicking the effect of a SHH organizer (SHH-organoid)^46^. SHH-organoids were grown and matured to 70 days, where they confirmed topography established at day 20 remained largely discrete and contained cell types of cortical, MGE, and hypothalamic lineages. Alternatively, microfluidic devices have been designed to generate defined exogenous chemical gradients by a planar dilution network to pattern the A-P axis of an entire neural tube ^47–50^. While such microfluidic gradient generator offers the adaptability of creating chemical gradients for a wide variety of signaling molecules in a controlled manner, it is still challenging to establish a steep gradient within tissues with low aspect ratios such as most organoids. In addition, microfluidic technologies have been applied mostly to adherent tissues, and the limited space in enclosed microfluidic channels prevents long-term culture or transferring of patterned tissues, which is essential for developing mature neural organoids. Moreover, to generate the chemical gradient in a microfluidic system, the use of peristaltic pumps and copious amounts of media are required, hindering its ability to scale up and be adopted by other research labs. Hence, a more accessible alternative method with the capability to reliably produce a gradient and allow for long-term culture is desired.

In the current study, we describe a novel strategy to generate <u>M</u>orphogen-gradient Induced <u>B</u>rain <u>O</u>rganoids (MIBOs). Leveraging the properties of passive diffusion to create a sustained chemical gradient of purmorphamine (Pur), a Shh agonist, we designed and optimized a microdevice for 3D tissue culture to induce D-V patterning within a single forebrain organoid. Using this technology, MIBOs were patterned for 19 days and further cultured out of the device long-term with observable maturation up to 5.5 months. Gradient quantification of 3.5-week-old MIBOs revealed robust gradient magnitudes for both NKX2.1 and PAX6. Single cell transcriptomic analysis of 4-month-old MIBOs revealed composition of cell types primarily arising from the LGE and CGE with contribution of the MGE to a smaller degree. Finally, MIBOs were functionally active as indicated by live calcium imaging analysis. Thus, MIBOs can be fine-tuned with specific morphogenic gradients to achieve spatially patterned and functionally mature brain organoids.

## Results

### Deriving MIBOs by an exogenous Pur gradient

We have previously developed a <u>L</u>ocalized <u>Pa</u>ssive <u>D</u>iffusion (LPaD) device to induce sustained concentration gradients of Pur to pattern 2D neuroepithelial cell sheets ^51^. In the current study, we have altered the design of LPaD for 3D organoid culture (Fig. S1). First, we added polydimethylsiloxane (PDMS) sidewalls to self-contain the culture medium. Doing so allowed us to reduce media usage by ten-fold. Further, to ensure that organoids are continuously exposed to the Pur gradient from the same direction, we embedded organoids in Matrigel to limit the rotation of the organoids (Fig. 1A, B). To facilitate the Matrigel embedding, we added a structure within the culturing area, allowing for the droplet formation needed to Matrigel embed our organoids at three varying distances from the Pur source (Fig. 1B).

**FIGURE 1.**
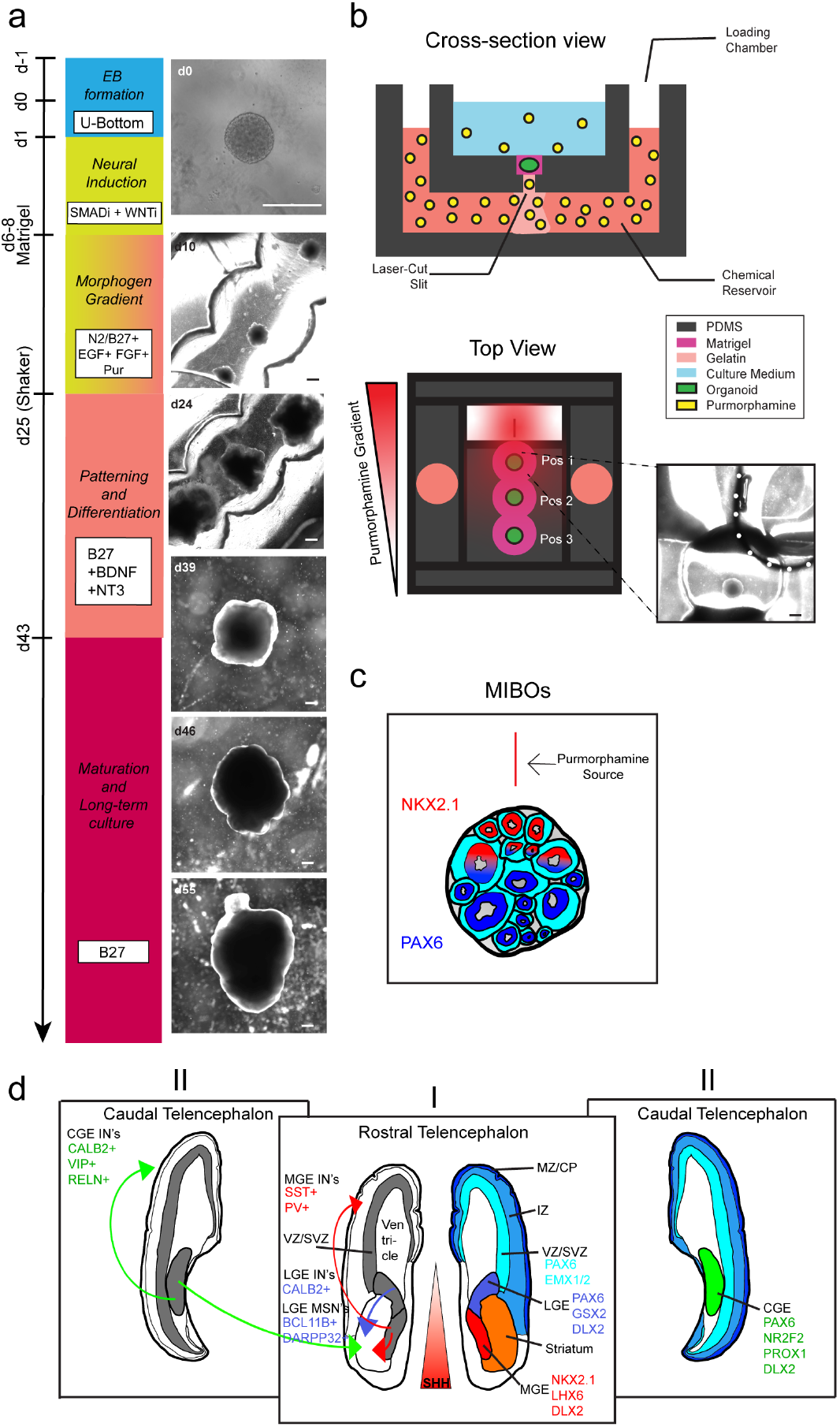
| Design and generation of MIBOs. (A) Schematic overview of culturing protocol in gradient device with representative bright field images. All scale bars are 500 µm. (B) Cross section of PDMS device showing the controlled passive diffusion of Pur molecules from the chemical reservoir into the culturing medium. Top view of PDMS device showing the three possible positions within the device and the effect of Pur gradient with distance from the source. (Dotted lines) Phase image of a freshly embedded organoid in a device with gelatin drop (white dotted line) over 200µm slit. (C) (Left) Cartoon representation of a patterned organoid in a PDMS device. Regions closest to the Pur source produce NKX2.1+ rosettes, while areas further away produce PAX6+ rosettes. (D) Cartoon renderings of coronal sections of the human telencephalon at 15 pcw made with references from the Allen Brain Atlas. The colored rostral telencephalon (Right) depicts MGE, LGE, and cortical regions of the embryonic brain along with markers expressed by the progenitors of each region. Black and gray rostral telencephalon depicts migratory routes populating the striatum and cortex with GABAergic projection neurons and INs from the LGE, CGE, and MGE along with common markers associated with each. Caudal telencephalon shows CGE, and markers expressed by its progenitors and INs.

We next cultured forebrain organoids using a modified version of the protocol developed by Cederquist et al., 2019 (Fig. 1A). Without activation of the SHH pathway, a default dorsal-anterior forebrain fate was anticipated. Once organoids were Matrigel embedded in our device, organoids were immobile without significant hydrogel degradation and were capable of being held in place for the subsequent 19 days in the device. Organoids continued to grow stably and exhibited neuroepithelial organization during this period despite immobilization (Fig. 1A). At day 25 (3.5 weeks), organoids were carefully removed from devices and transferred into ultra-low attachment dishes which were then placed on orbital shakers for the remainder of the culturing scheme. Long-term organoids grew to maturity for further analysis using immunohistochemistry (IHC), single cell RNA sequencing, and calcium imaging. Using this two-step protocol, MIBOs cultured in our devices are not limited by the space constraints of the device and can be cultured long-term for further maturation and characterization after the initial patterning period.

### Successful Dorsal-Ventral patterning in MIBOs

Having established organoid viability using our device, we next sought to characterize MIBOs that had been exposed to a concentration gradient of 1µM Pur. Here, we observed that MIBOs had simultaneous but mutually exclusive expression of the dorsal (PAX6) and ventral (NKX2.1) forebrain markers (Fig. 2A). With 1µM Pur, we robustly observed NKX2.1-expressing as well as PAX6-expressing neural rosettes within a single organoid (Fig. S2). This unique expression pattern further emphasizes the ability for our device to procure ventral gradation even within a single progenitor domain. Next, we developed a MATLAB code (see Methods) that uses a sobel gradient operator to quantify the gradient magnitude of a grayscale image of PAX6 or NKX2.1 staining (Fig. 2B, C). The analysis showed a significant increase in the gradient magnitude for MIBOs compared to our positive controls (adding 1µM Pur directly into the culture medium) and negative controls (no Pur added). As expected, the positive control showed a high degree of NKX2.1 expression in organoids (Fig. 2A, C). In contrast, the negative control exhibited homogenous PAX6 expression throughout the organoid (Fig. 2A, C).

**FIGURE 2.**
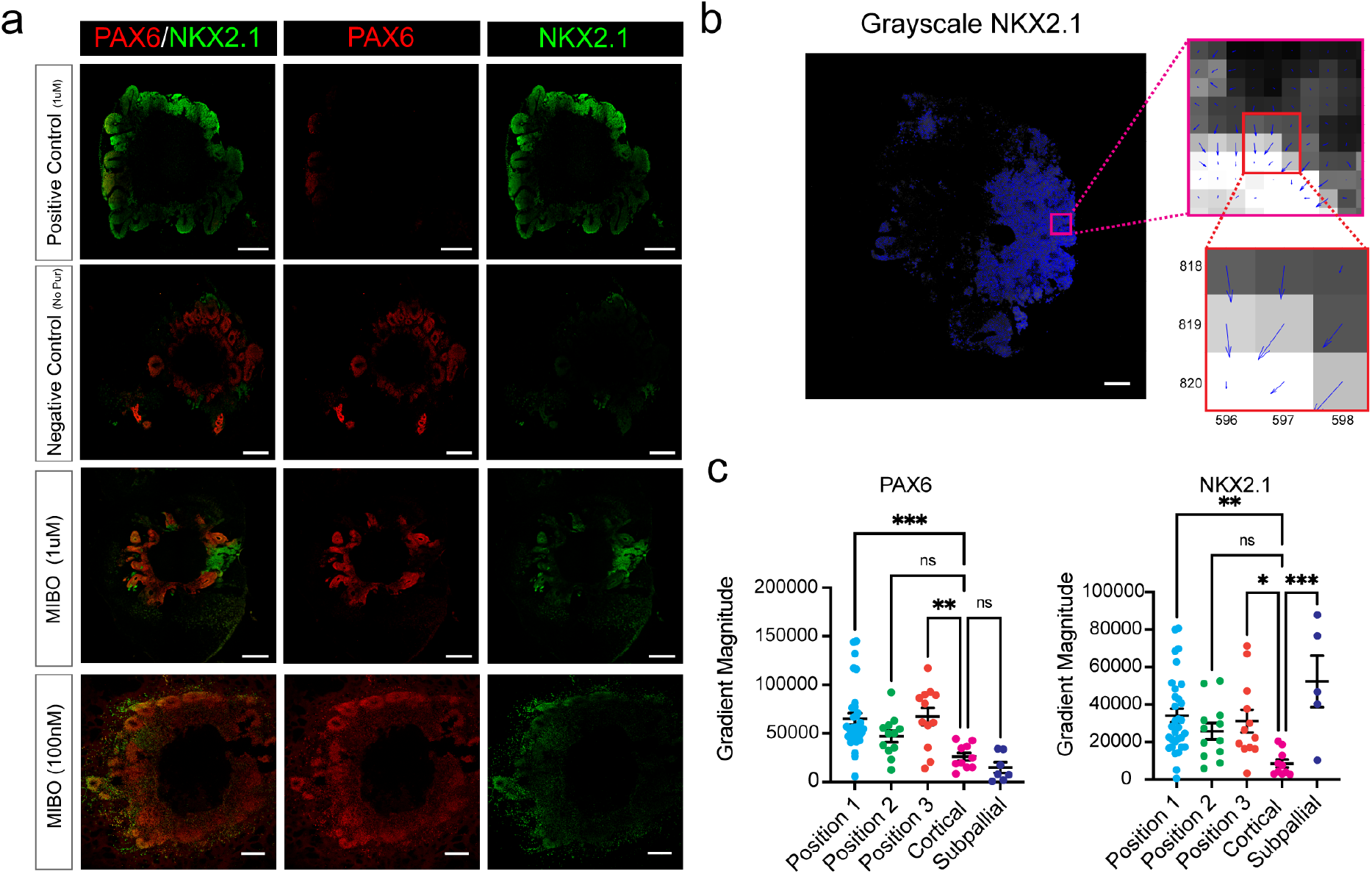
| Characterizing the patterning effects of Pur gradient on forebrain organoids. (A) Comparison of brain organoids grown in PDMS devices and subjected to varying concentrations of Pur with positive (adding 1µM Pur directly into the culture medium) and negative controls (no Pur added). All scale bars are 500 µm. Scale bar for 100nM condition has an 200µm scale bar. (B) (Left) MATLAB analysis of grayscale NKX2.1 gradient magnitude. (Top) Zoomed in image showing blue quivers for each pixel. Size of quiver corresponds to size of gradient magnitude and quiver angle displays the direction of greatest increase in pixel intensity. (Bottom) 3x3 neighborhood of pixels illustrating how the center pixel establishes a gradient direction and gradient magnitude based off its neighbors. (C) (Left) Gradient magnitude measured in sections stained for PAX6 across all conditions. (Right) Gradient magnitude measured in sections stained for NKX2.1 across all conditions. Error bars represent +/- S.E.M. Statistical significance was obtained by one-way ANOVA with Dunnett’s test for multiple comparisons. *p<0.05, **p<0.01, ***p<0.001.

To determine if the cell fate patterning in MIBOs depends on the source concentration of Pur, we tested the effects of adding a 100µM or 100nM Pur concentration to the chemical reservoir. These two concentrations had originally been used for varying degrees of patterning effects in the 2D culture system ^51^. We observed full ventralization of the organoids exposed to a 100µM Pur gradient (Fig. S2C). NKX2.1 was uniformly expressed throughout these organoids. Interestingly, at 100nM condition, we found that PAX6 was largely expressed by the radially arranged progenitors of the rosette structures. On the other hand, NKX2.1 expression was still detected, albeit minimally scattered across the regions just distal to the rosettes (Fig. 2A).

Finally, we examined the effect of distance from Pur source on D-V patterning. We designed the Matrigel embedding area with three equally ascending distances from the Pur source, which we have termed here, Position 1, Position 2, and Position 3 (Fig. 1B). Position 1 showed the most consistency in producing D-V patterning within the forebrain organoids. Within Position 1, we observed consistent generation of regions expressing NKX2.1 within VZ progenitors, indicating emergence of MGE-like regions from the side of the organoid exposed to the highest concentration of Pur (Fig. 2A). The rest of the organoid expressed PAX6, indicating a more dorsal fate. While Position 2 and 3 organoids occasionally showed similar D-V patterning, the PAX6 and NKX2.1 expression was deemed stochastic (Fig. S2A). As a result, hereafter we focused our efforts on further characterizing the MIBOs produced in Position 1 induced by 1 µM Pur source.

### Investigating cell fates arising from progenitors of MIBOs through single cell RNAseq

Cederquist et al. had previously documented a stark difference in the size of PAX6+ versus NKX2.1+ expressing rosettes in the SHH organoids, where NKX2.1+ rosettes tended to be elongated when compared to the more circular PAX6+ rosettes. Within our patterned organoids, at the border of where markers transition, we noted several rosettes that partially expressed NKX2.1 and PAX6 (Fig. S2B). Additionally, many PAX6-expressing rosettes also showed elongated morphologies, prompting us to further investigate the fate of mature cells produced by these patterned organoids in single cells.

Single cell transcriptomic analysis of 4-month-old MIBOs revealed a lack of cortical or excitatory markers within our sample (Fig. 3). Instead, we found that a large majority of our cells were subpallial in nature. Specifically, cluster cell annotation based on canonical markers identified LGE- and CGE-derived GABAergic cells with a small group of MGE-derived cells (Fig. 3A-C, Fig. S3A-C, Table S1). From this data we reasoned that PAX6 staining pattern observed at 3.5 weeks specifically marked dorsal subpallial progenitors of the LGE/CGE and not cortical progenitors of the pallium ^11, 19^. Therefore, the patterning created in MIBOs was more representative of the D-V patterning observed in the GEs.

**FIGURE 3.**
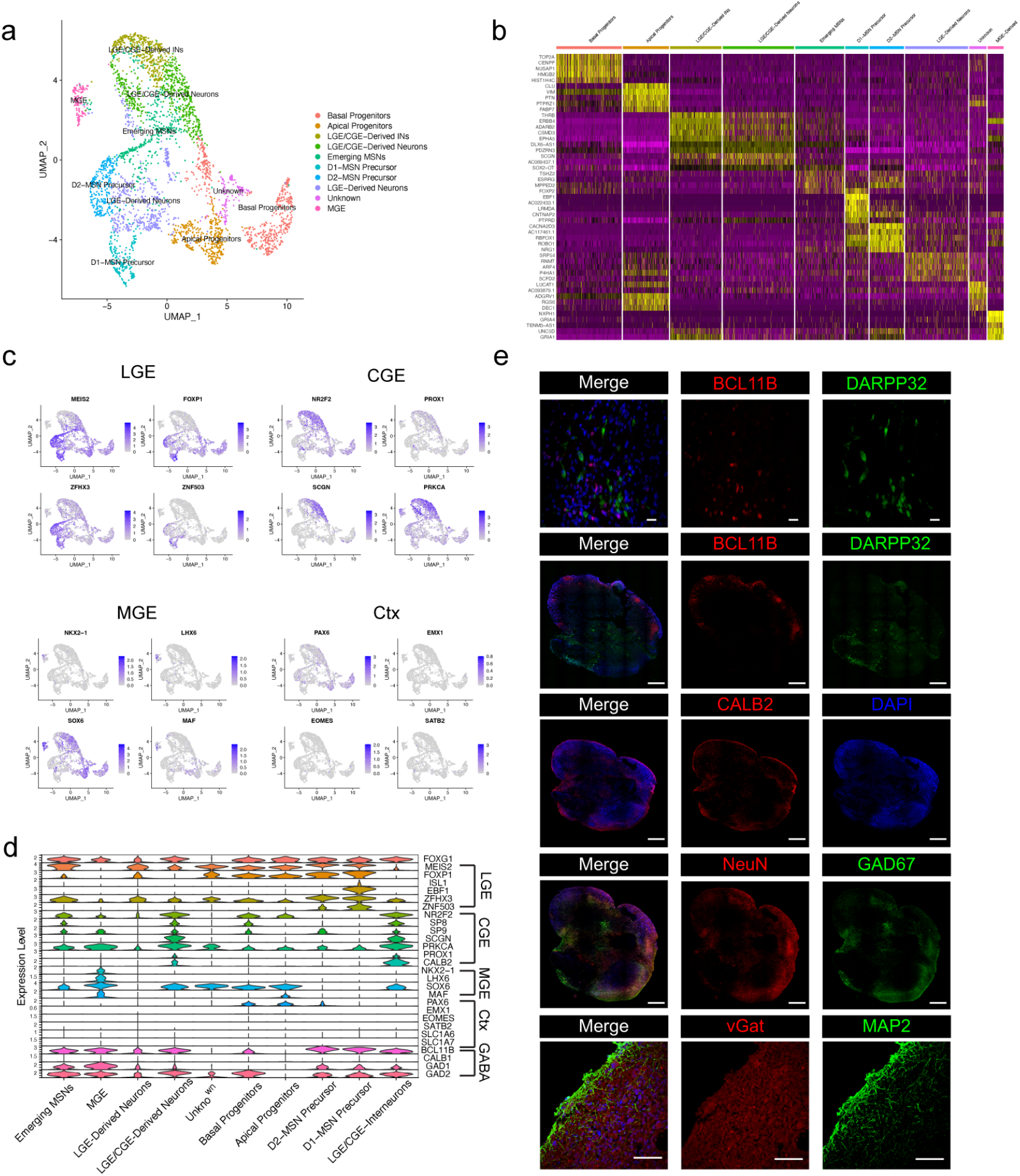
| Single cell transcriptomics reveals GE fates in MIBOs. (A) UMAP embedding of 3,366 cells dissociated from Position 1 MIBOs at 4 months. (B) Heatmap showing the top 10 genes expressed in each annotated cluster. (C) Feature maps showing expression for the top marker genes in the LGE, CGE, MGE, and Cortex. (D) Violin plot highlighting representative lineage marker genes across annotated cell clusters. (E) Representative confocal images from 3.5-month-old organoid sections immunostained with specific markers labeled. Scale bar for 40x BCL11B is at 50 µm. Whole organoids stitched images use an 500µm scale bar. MAP2 and VGAT was captured at 60x and used an 100µm scale bar.

Indeed, cell annotations of our single cell data found evidence of LGE-derived clusters with robust expression of known LGE-markers, including MEIS2, FOXP1, and EBF1 (Fig. 3C, D). Additionally, emerging medium spiny neuron (MSN) and MSN precursors for D1 and D2 subtypes were identified within our LGE-derived clusters (Fig. 3A-D) ^21^. The GABAergic projection neurons of the LGE are thought to be primarily produced in the ventral LGE (vLGE), with GABAergic INs more heavily produced by the dorsal (dLGE) ^11, 52, 53^. Interestingly, the GABAergic IN population in our sample was almost entirely CALB2+ cells. Some of these CALB2+ INs could be LGE derived while others derive from CGE progenitors due to high coexpression with NR2F2, SP8, PROX1, and PRKCA ^20, 21, 54, 55^ (Fig. 3C). The LGE/CGE-derived neurons cluster was identified by coexpression of LGE/CGE markers with the presence of BCL11B+ and SOX6+ (Fig. 3D, S3E). BCL11B is found with high fidelity in mature MSNs, as well as early postmitotic MSNs ^56^. Moreover, while SOX6 is strongly expressed in the MGE and CGE mantle zone (MZ), the MZ of the LGE (FOXP1+) lacks SOX6 expression and is notably the region of MSN maturation ^56, 57^. Finally, MGE derived neurons were identified in a small cluster that expressed the canonical markers NKX2.1, LHX6, and SOX6 (Fig. 3C, D).

We next sought to validate our results by immunostaining MIBOs that had been matured and collected at 3.5 months. Again, we saw wide expression of BCL11B+ and even some DARPP32+ cells indicating the presence of GABAergic projection neurons, and possibly emerging MSNs (Fig. 3E). The most abundant INs in MIBOs were CABL2+ cells that seemed to be undergoing tangential migration with most cells located at the outermost region of the organoid ^58, 59^ (Fig. 3E, S3G). While many cells within our MIBOs appear to be undergoing migration, a significant portion of them were differentiated neurons expressing mature markers for GABAergic subtypes, such as RBFOX3, GAD1, and SLC32A1 (Fig. 3E, S3F).

In summary, we have shown that MIBOs can produce a wide variety of cells derived from the GEs. At the current concentration of 1µM Pur, NKX2.1+ regions of MIBOs are exclusive to small sections that continue to produce MGE derived cells into maturity. The rest of the MIBOs represents the D-V axis of the LGE and CGE, thus producing a range of GABAergic projection neurons and primarily CALB2+ INs. These results indicate that PAX6 expression and rosette morphology at 3.5 weeks were likely representative of dorsal LGE/CGE apical progenitors.

### Characterizing the functioning neural circuits in MIBOs using calcium imaging analysis

To better characterize the neural activities within MIBOs, we performed live Ca^2+^ imaging in MIBOs and a subpallial control organoid at 5.5 months (Fig. 4A). We observed stable and consistent levels of neural activity in their spontaneous Ca^2+^ spikes, spike amplitudes, and synchronicity of spikes (Fig. 4B, C). To further investigate if cell fate patterning influences neural maturation, Ca^2+^ activity in MIBOs across three positions was comparable to that of control subpallial organoids (Fig. 4D). Though we observed a minor position variability in their neural activity, all three positions showed robust synchronous and spontaneous firing with expected metrics as previously published ^60^. Hence, our results show free floating MIBOs that were cultured long-term out of the device yield active and viable organoids with mature neural network characteristics.

**FIGURE 4.**
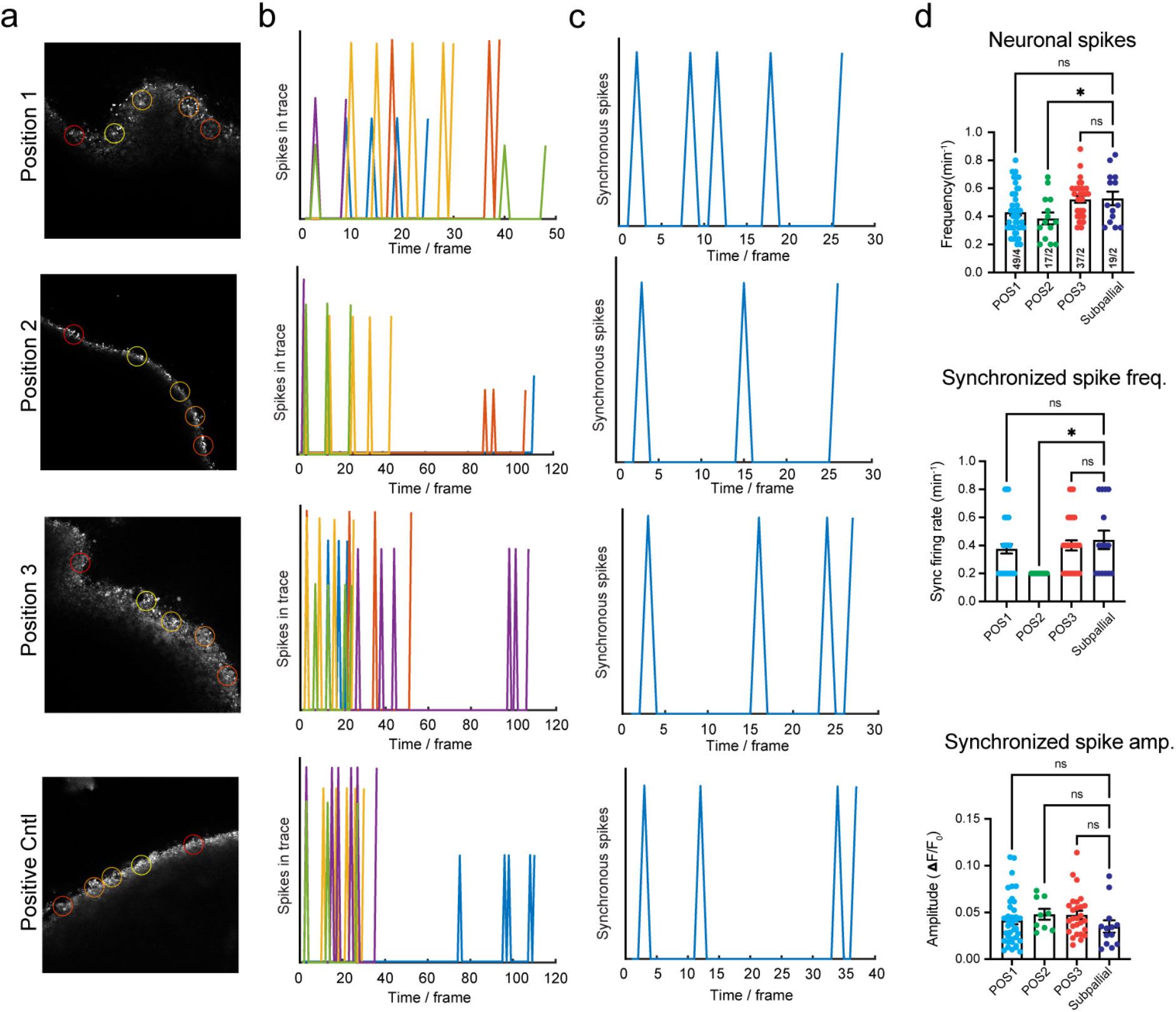
| Neuronal activity captured through calcium imaging of MIBOs grown long-term. (A) Representative confocal images of intact brain organoids during live Ca2+ imaging using X-Rhod-1 dye. Colored circles represent regions of interest (ROIs) selected for analysis. (B) Corresponding spike activities captured within ROIs during recording. (C) Network activity detected in brain organoids by synchronous spiking across multiple ROIs. (D) Averaged data for synchronous spikes detected in each condition, as well as amplitudes (dF/F0) and frequencies of spontaneous spike activity, are shown in scatter plots. Data points represent averaged data from a single field of view (FOV) consisting of 5 ROIs per FOV. At least 5-7 FOVs are taken from each organoid and 2-4 organoids were used per experiment. Error bars represent +/- S.E.M. Statistical significance was obtained by one-way ANOVA with Dunnett’s test for multiple comparisons. *p<0.05.

## Discussion

Developmental biology studies strongly support the diffusion and reaction-diffusion of morphogens in inducing tissue patterning. The D-V patterning in telencephalon is believed to be induced by Shh produced by the notochord and floor plate, as well as BMP produced in the roof plate. However, the D-V patterning mechanism in the GEs is still poorly defined. Here, we demonstrated that exposing forebrain organoids in a gradient of Pur is sufficient to generate GEs with segregated MGE region and CGE/LGE region. Compared with microfluidics-based approaches, our device takes advantage of passive diffusion of chemicals to generate a sustained chemical gradient without the need for continuously pumping and significantly reduced media consumption. Our open-chamber design allows for convenient organoid retrieval and extended culture, which is usually challenging in enclosed microfluidic devices.

Typically, in both 2D and 3D systems, the use of dual SMAD inhibition with WNT inhibition is thought to produce a primarily cortical phenotype ^44, 61, 62^. Similarly, in chimeric SHH-organoids, the PAX6+ regions also represented cortical fate ^46^. Surprisingly, PAX6 expression observed in 3.5-week-old MIBOs was primarily identifying the dorsal GE regions of the LGE and CGE. However, given that MIBOs were exposed to a Pur gradient without antiparallel source of Shh antagonism (e.g., BMP4), we speculate that the low levels of persistent Pur throughout the media was sufficient to push a dorsal subpallial fate in the regions furthest from the Pur source. In MIBOs, Pur gradient begins at around 1 week, immediately after the establishment of forebrain neural stem cell lineage in the EBs. Namely, the ‘temporal adaptation’ model suggests that, initially, cells will be highly responsive to Shh stimulation ^63, 64^. Furthermore, protocols attempting to produce LGE derived cell types will typically use lower concentrations of Pur, ranging from 0.5 to 0.65 µM ^39, 65–68^. Therefore, it is likely that the Pur gradient created by our MIBO device was more conducive for the generation of subpallial tissues in general. Excitingly, the device platform is promising for testing and characterizing the manipulation of different Pur concentrations in such a way that different concentration effects could be used to establish new protocols with varying degrees of MGE to LGE/CGE ratios. For example, we predict that an increase in Pur concentration in our chemical reservoir would likely generate increasingly larger MGE regions within our organoid. Potentially, Pur could be increased to such a degree that the D-V patterning of the MGE itself is induced, allowing closer examination of the development of dorsal MGE SST+ and ventral MGE PV+ INs ^11, 69–71^.

Our single cell transcriptomic analysis of MIBOs revealed a fascinating diversity of subpallial derived cell types. Specifically, we identified a small cluster of MGE derived cells in addition to vLGE-derived MSNs and dLGE/CGE derived INs. Cell types were annotated using key marker genes that have been well documented for each region ^11, 19–21, 54, 57, 71–78^. While the markers for delineating MGE-derived cells have been established and are unique to the region, the LGE and CGE tend to share many of the same markers (i.e. NR2F2, PAX6, MEIS2) with differences in expression levels serving as the only indicator of regional identity ^55, 79^. Accurately discerning the LGE from the CGE still remains a challenge for the field. Traditionally, the only way to definitively discriminate the regions of the GEs were by anatomical landmarks (bulges). Recently, the field has made great strides that could help improve our understanding of these GEs through the use of single cell RNA-sequencing on rodent and human embryonic tissues to further identify novel unique markers for each region ^20, 21, 75, 78^. Due to this ambiguity, it is possible that some clusters in our dataset house cells from both the LGE and CGE jointly. For example, the dLGE is known to produce CALB2+ INs that are fated for the olfactory bulb, migrating through the rostral migratory stream ^4, 16, 71, 80^. Likewise, the CGE produces CABL2+ INs that will tangentially migrate towards the cortex and/or striatum ^9, 81^. Due to the similarity of gene expression between both regions, it is possible that the CALB2+ INs of both the LGE and CGE are clustered together in our dataset.

Interestingly, the majority of INs found in our MIBOs were CALB2+ cells. However, the CGE has been well characterized in its ability to produce CALB2+, VIP+, and RELN+ INs ^17, 82^. This begs the question as to why our organoids primarily produced CALB2+ INs. While we did identify a small number of VIP+ expressing GABAergic cells in our LGE/CGE cluster (Fig. S3H), it is plausible that we sequenced MIBOs at an age when VIP IN production was just beginning. Developmentally, the MGE produces cortical INs at an earlier time point than the CGE ^17, 83^. MGE-like subpallial organoids were reported to begin IN production at around Day 50 for SST+ and Day 200 for PV+ ^28^. Therefore, it is possible that allowing MIBOs to mature beyond 4 months would yield a higher amount of VIP+ or RELN+ INs. Further, CGE-derived INs were reported not to express their subtype markers until they have finished their migration and settled in their final destination, usually within the outermost layers of the cortex ^17, 84–86^. It is theorized that, post migration, exogenous signals and local electrophysiological input from the local environment triggers mature marker expression. VIP+ cells often co-express CALB2+ as well. Therefore, there is potential for a portion of the currently identified migrating CALB2+ INs to mature into VIP+ or RELN+ co-expressing INs. One additional clue for CGE tissue production in MIBOs is the presence of CALB1+ INs stained for and visualized in 3.5-month sections (Fig. S4). Finally, we must consider the temporal effects on IN production in the CGE. Previously, it has been shown that the MGE alters its ratio of SST+ and PV+ IN production temporally ^83^. In contrast, the IN ratio of VIP+, CR+, and RELN+ produced by the CGE remains roughly unchanged throughout embryonic maturation further hinting at the possibility of even distribution of INs if MIBOs are allowed to mature beyond 4 months ^17^.

Though we have focused on the ability of our devices to D-V pattern the subpallium, ultimately, our device also has the potential to be a platform that can be further modified to produce more complex neural structures. One option is to make use of temporal adaptation by introducing the Pur gradient at a later time point; once cortical fates have been established and sensitivity to Shh has waned. Doing so might generate pallial and subpallial tissues within a single organoid using a single morphogen. Alternatively, this could also be achieved with an antiparallel gradient of morphogens mimicking BMP and Wnt. In the current iteration of our device, we used a single source of morphogen for the ventralization of our tissues. Future modifications could introduce an additional source that releases a dorsalizing small molecule such as cyclopamine (Shh antagonist) or BMP4. We predict that the exposure to the opposing gradients could create MIBOs that express both subpallial and cortical markers. An organoid model that intrinsically produces subpallial and cortical features could be extremely useful for teasing apart the questions surrounding embryonic neurodevelopment and potentially provide a useful model for studying the effects of neurodevelopmental disorders on IN production and migration.

In conclusion, in this work, we report the fabrication of an easy-to-use PDMS device to generate a sustained chemical gradient for the organoid patterning. We showed that Pur gradient could induce MIBOs mimicking the D-V patterning observed in the GEs. Single cell transcriptomic analysis revealed a rich cell diversity of GE derived GABAergic subtypes, including dorsally derived CALB2+ INs, vLGE MSN precursors, and a small population of MGE derived cells. Live Ca2+ imaging analysis showed that MIBOs matured into intact organoid structures with functional neurocircuitry. Our system, therefore, could provide a new strategy for generating properly patterned and regionalized neural organoids, a vital foundation for modeling neurodevelopment and disease states.

### Limitations of the study

Our current version of MIBO device fabrication requires multiple PDMS slabs being cut out, carefully assembled and sealed, which can be time consuming. However, since a majority of the device is made of PDMS, 3D printing molds and consolidating key pieces for a PDMS cast of the device would greatly decrease assembly time. In this study, we have fully relied on NKX2.1 staining to reliably indicate the region of the organoid closest to the Pur source (laser-cut slit). To further validate the direction of the gradient, a polymer microbead could be manually embedded to the side of the organoid closest to the Pur source. Confirmation would require colocalization of NKX2.1 expression on the same side as the microbead. Additionally, while we observed reproducible patterning in a single human embryonic stem cell line (WA-01), reproducibility and robustness of the technology could be tested with additional ESC and induced Pluripotent Stem Cell (iPSC) lines.

## Acknowledgements

This work was supported by NIMH (R01 MH122519 to C.P., R21 MH130843 to Y.S. and C.P.), UMass IALS/BMB faculty start up fund (to C.P.), Tourette Association of America (Young investigator award to C.P.), UMass IONS Seed funding, and NIGMS T32 BTP training program (T32 GM135096 to N.P.). We would like to thank members of the Sun and Pak labs for critical feedback and acknowledge the support from UMass IALS Light Microscopy Core and Genomics Core.

## Author contributions

The study was designed by Y.S., C.P., and N.P. N.P. executed all experiments and analysis with the help of K.D., F.Y., R.S., B.M-M., and R.R. Y.S., C.P., and N.P. interpreted the data, generated figures, and wrote the manuscript.

## Declaration of interests

The authors listed (Yubing Sun, ChangHui Pak, Feiyu Yang, and Narciso Pavon) have filed a provisionary patent related to this work (Docket Number: UMA 23-023, Title: “A chemical gradient inducing device and methods for developing patterned organoids.”).

## STAR Methods

### Key resources table

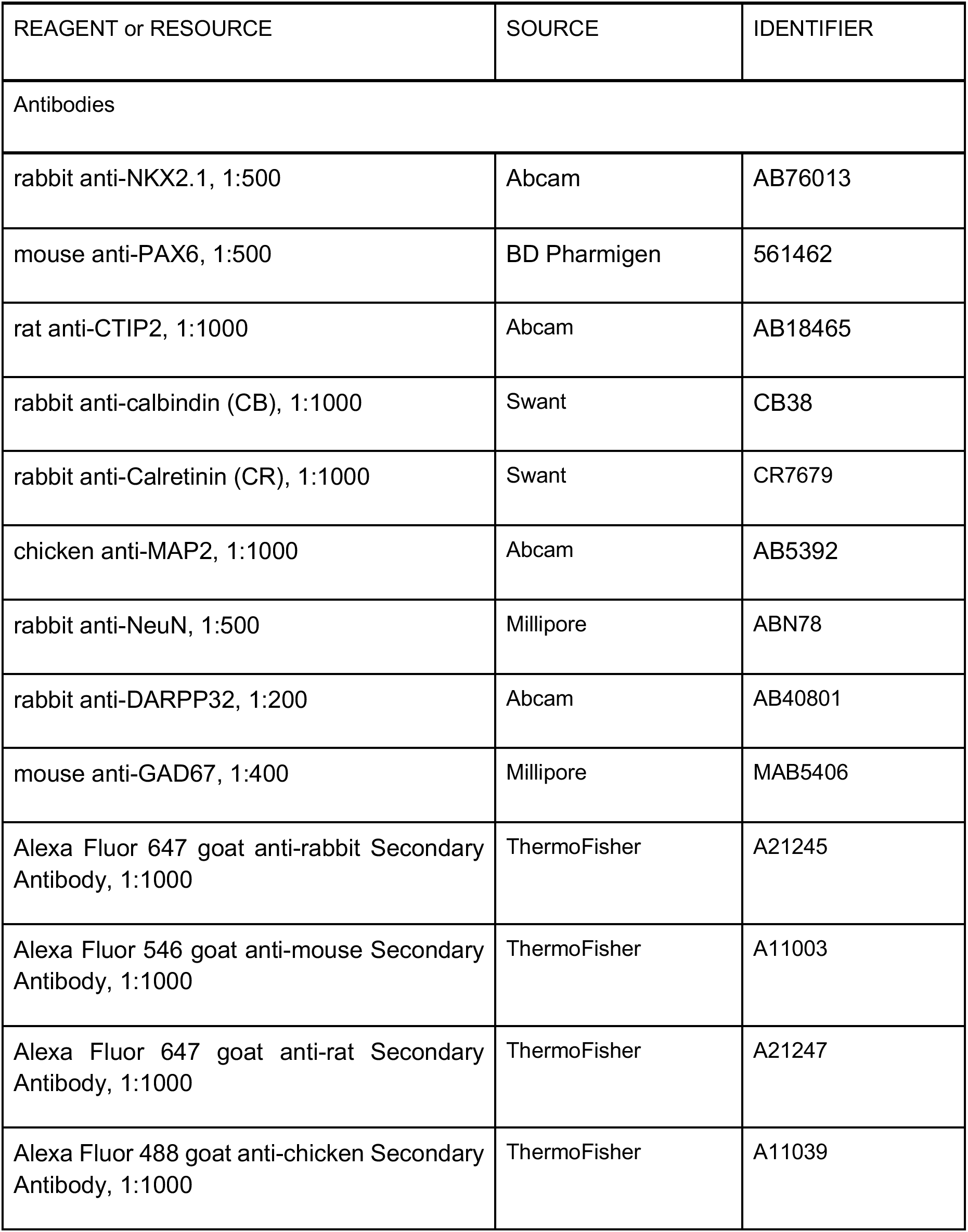
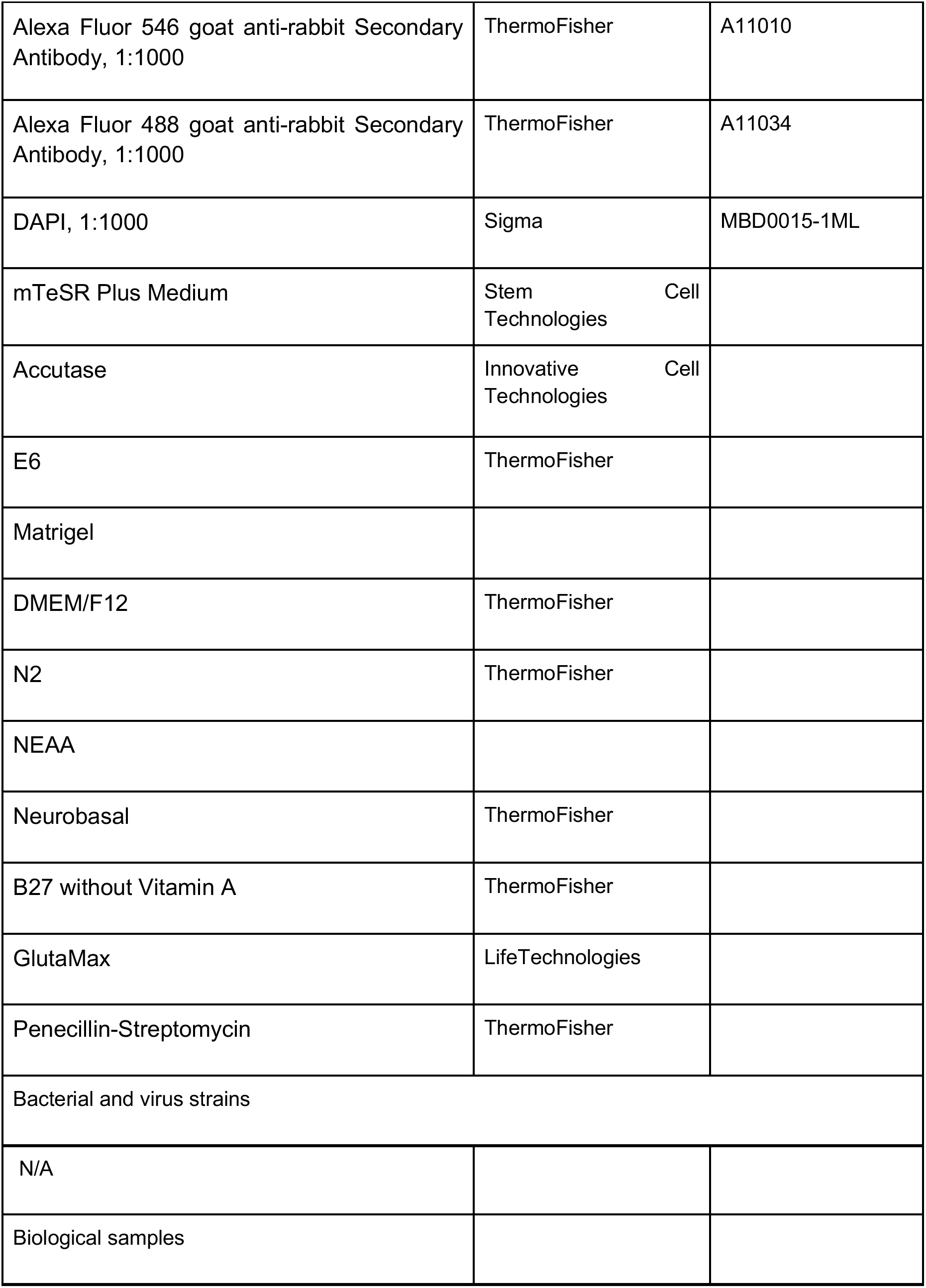
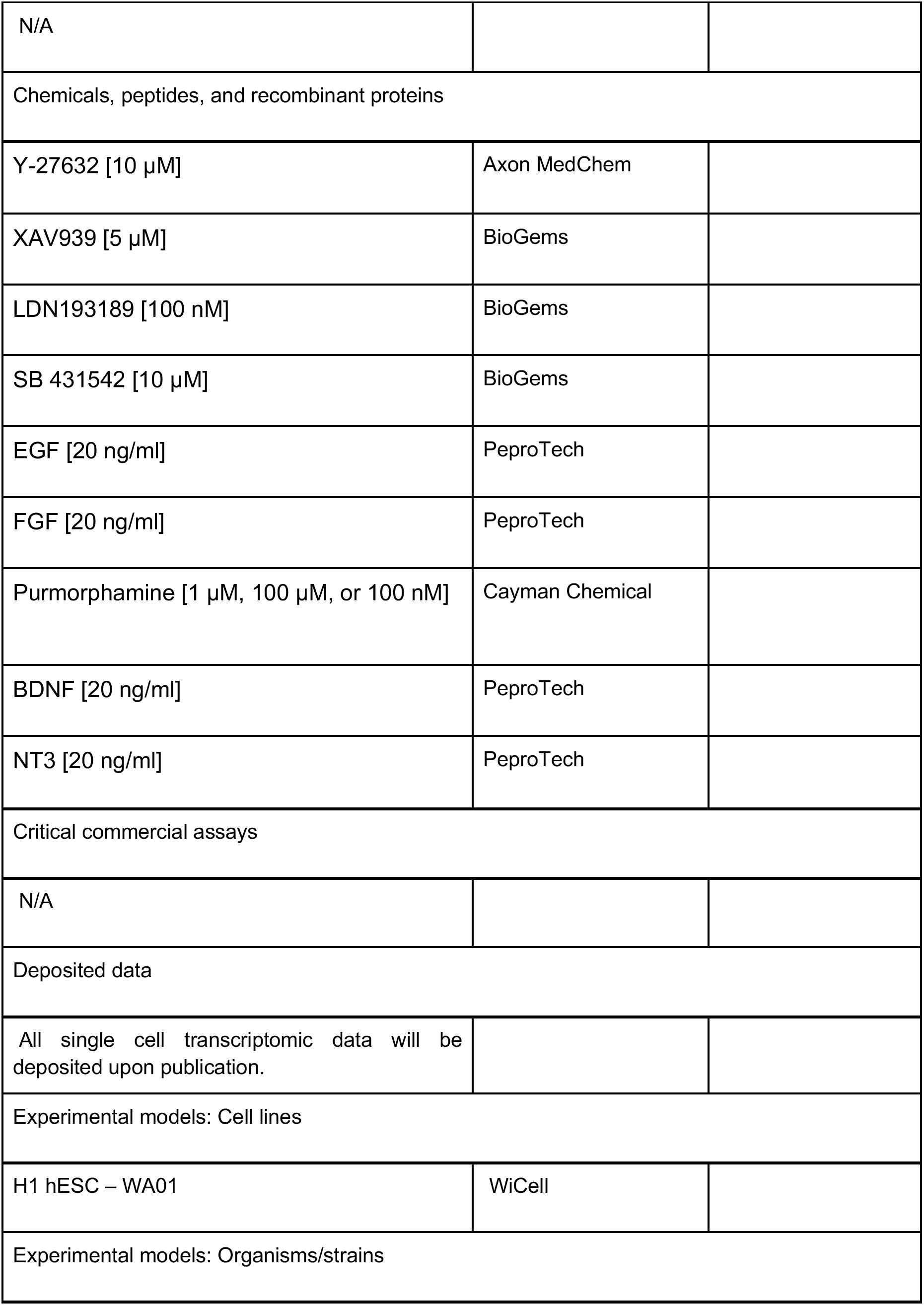
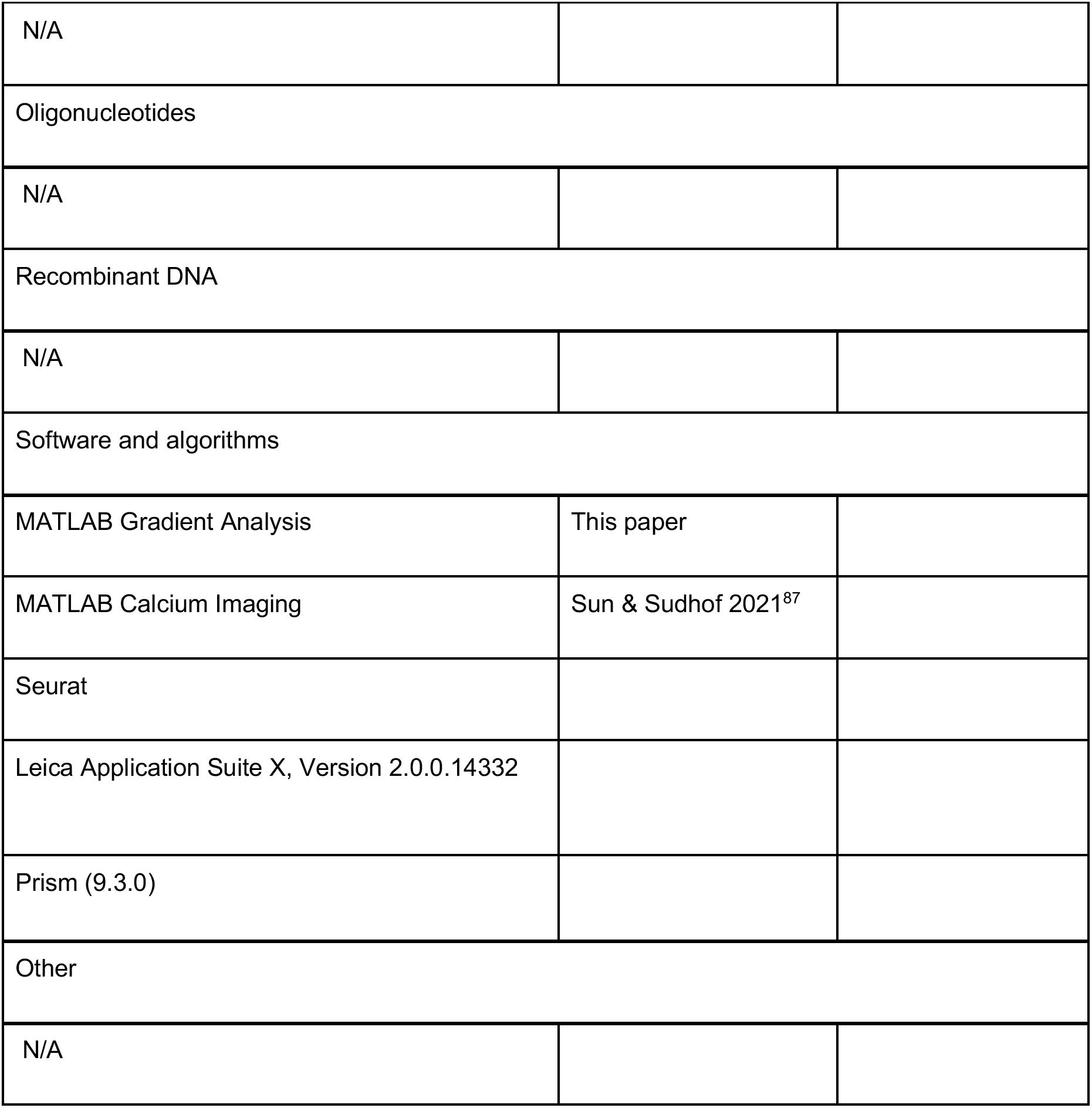

### Experimental models and subject details

#### Cell culture

H1 hESC line were grown in a standard 6-well plate with a 9.6 cm^2 culturing surface area coated with Matrigel diluted in DPBS and maintained with mTeSR Plus medium (Stem Cell Technologies) in feeder-free conditions. Y-27632 [10 µM] (Axon MedChem) was added to medium for all cell passaging. Cells were routinely tested for mycoplasma.

#### MIBOs generation

First, hPSCs were dissociated into single cells using Accutase (Innovative Cell Technologies). We then seeded these cells at a density of 10k cells per well in an Ultra-Low Attachment 96 Well, Round Bottom plate (Costar). Cells are allowed to aggregate overnight in mTeSR Plus with Y-27632 [10 µM] and XAV939 [5 µM] (BioGems). After 24 hours, EB cell aggregates will form, and culture medium will switch to E6 (ThermoFisher) containing LDN193189 [100 nM] (BioGems), SB 431542 [10 µM] (BioGems), and XAV939 [5 µM] (BioGems) for 8 days during neural induction. On the eighth day of neural induction, three organoids are transferred into MIBOs device using 200uL wide bore pipette tip (Thermo Scientific), with excess media removed using standard bore pipette tip. 30uL of ice cold Matrigel is promptly added for each organoid in a MIBOs device, and the pipette tip is used to center each organoid within the device embedding areas. Once embedded, culture medium is switched to a neural differentiation medium consisting of a 1:1 mixture with a base of DMEM/F12 (ThermoFisher) with N2 (ThermoFisher) and NEAA (ThermoFisher) and a base of Neurobasal Medium (ThermoFisher) containing B27 without Vitamin A (ThermoFisher), GlutaMax (LifeTechnologies), and Penicillin-Streptomycin (ThermoFisher). To this 1:1 base mixture we added EGF [20 ng/ml] (PeproTech), and FGF [20 ng/ml] (PeproTech). For the chemical reservoir medium, the same medium was used with the addition of varying concentrations of Purmorphamine [1 µM, 100 µM, or 100 nM] (Cayman Chemical). At Day 12, MIBOs are weaned off N2 by decreasing concentration from 1:100 to 1:200. On Day 25, MIBOs were carefully removed by gently dislodging Matrigel droplets using a standard bore 200uL pipette tip, and promptly transferred into ultra-low attachment 10 cm petri dishes using 1000uL wide bore pipette tips (Genesee Scientific). At Day 25, MIBOs were placed on an orbital shaker and cultured in medium containing only Neurobasal, B27 without vitamin A, Glutamax, Penicillin-Streptomycin, BDNF [20 ng/ml] (Peprotech), NT3 [20 ng/ml] (Peprotech). For long term culture, at day 43, all morphogens were removed and MIBOs culturing continued with Neurobasal, B27 without vitamin A, Glutamax, and Penicillin-Streptomycin alone.

#### Cryopreservation and sectioning

Organoid samples were collected at day 25 and day 101. Collected organoids were then washed 3 times with PBS and fixed with 4%Paraformaldahyde for 24 hours in 4 °C. After fixing is complete, samples are washed with PBS and stored in 30% sucrose solution for 24-48 hours in 4 °C. Organoids are encased in gelatin solution (10%gelatin with sucrose/PBS) followed by the flash freezing process using a mixture of dry ice and ethanol. Gelatin blocks are then stored in -80 °C for long term storage and eventually cryosectioned between 12 to 25 micron section thickness. Sections directly adhere to SuperFrost Plus slides (Fisher) and promptly used for immunohistochemistry. Unused slides are stored in -20 °C.

#### Immunostaining

Sections are washed three times using 0.2% Triton-X in PBS (0.2%PBS/T) and blocked for 1 hour at room temperature (RT) using 10% normal goat serum in 0.2%PBS/T (blocking solution). After 1 hour, sections receive a quick rinse with 0.2%PBS/T and are incubated overnight at 4 °C with blocking solution containing diluted antibodies. The following day slides are washed three times with 0.2%PBS/T and incubated with secondary antibodies and DAPI diluted in blocking solution for 2 hours at RT. After the incubation period, slides are washed three times with 0.2%PBS/T and promptly mounted using Fluoromount mounting media (Southern Biotech). Primary antibodies used are as follows: rabbit anti-NKX2.1 [Abcam, AB76013; 1:500], mouse anti-PAX6 [BD Pharmigen, 561462; 1:500], rat anti-CTIP2 [Abcam, AB18465; 1:1000], mouse anti-SATB2 [Abcam, ab51502; 1:400], rabbit anti-calbindin (CB) [Swant, CB38; 1:1000], chicken anti-MAP2 [Abcam, AB5392; 1:1000], rabbit anti-Calretinin (CR) [Swant, CR7679; 1:500], rabbit anti-NeuN [Milipore, ABN78; 1:500], rabbit anti-DARPP32 [Abcam, ab40801; 1:200], and mouse anti-GAD67 [Milipore, MAB5406; 1:400]. Secondary antibodies used: Alexa Fluor 647 goat anti-rabbit [ThermoFisher, A21245, 1:1000], Alexa Fluor 546 goat anti-mouse [ThermoFisher, A11003, 1:1000], Alexa Fluor 647 goat anti-rat [ThermoFisher, A21247, 1:1000], Alexa Fluor 488 goat anti-chicken [ThermoFisher, A11039, 1:1000], DAPI [Sigma, MBD0015-1ML; 1:1000], Alexa Fluor 546 goat anti-rabbit [ThermoFisher, A11010; 1:1000], Alexa Fluor 488 goat anti-rabbit [ThermoFisher, A11034; 1:1000].

#### Image Acquisition

Images of sectioned organoids were imaged using a Leica DMi8 microscope with LAS X Software (Leica Application Suite X, Version 2.0.0.14332). A 10x objective was used to obtain multiple images of a single organoid and then merged to create a large image containing a full organoid section (Fig 2, S2, Fig 3). 60x, images were captured using Nikon A1R25 confocal microscope.

#### Calcium Imaging

Calcium dye mixture is prepared with 1mM X-Rhod-1 AM dye (Invitrogen) diluted in modified HEPES buffer (130mM NaCl, 5mM KCL, 2mM CaCl2, 1mM MgCl2, 10mM HEPES, 10mM Glucose, ∼pH 7.4 adjusted with NaOH). Organoids are then incubated in calcium dye mixture for 15 minutes at RT. After incubation period is over, organoids are quickly washed with the modified HEPES buffer once and imaged using a confocal microscope (Nikon, A1R25). A glass bottom petri dish (MatTek) was used for all imaging and a stable temperature of 37 °C was maintained using an Ibidi stage heater. Time lapse images are captured at 250 ms intervals for a period of 5 mins.

#### Calcium Imaging Analysis

Raw images are extracted using NIS Elements and analyzed using a stimulation-free MATLAB protocol as demonstrated previously (Sun &Sudhof). Using the MATLAB protocol, we first create a Maximum Intensity Projection (MIP) by ‘stacking’ the time lapse images captured using the confocal microscope. The MIP serves as a guide for selecting 5 active regions of interests (ROIs). Each ROI was contained to a 50 μm diameter. A time interval was determined by taking the recording duration (in seconds) over total frames i.e for our 5-minute recordings we used 300/109 to obtain a time interval of 2.75. The analysis tracks the changes in pixel intensity within ROIs for one field of view (FOV). The recorded changes in intensity for one FOV are used to obtain a frequency of spiking activity and instances where spiking activity occurs across more than one ROI are used to obtain synchronous spikes and a synchronous firing rate. Amplitude is obtained by using the mean value of DF/F0 from individual peaks.

#### Live Single Cell Dissociation

3.5-month-old organoids were transferred to a petri dish and rinsed with HBSS (1 bottle HBS salt, Sigma H2387-10X1L; 1 mL 1M HEPES, pH 7.3; 0.35g NaHCO3; 100 mL autoclaved ddH2O). After rinsing, organoids were cut into smaller pieces using a sterile razor blade. A small amount of Digestion Solution (1900ul HBSS (-/-), 50ul papain, 11 uL of 0.5M EDTA, 7 mg L-Cysteine) was added to the petri dish and a wide bore tip was used to transfer the smaller pieces into the Digestion Solution that was sterile filtered with 0.22 µm. The solution was incubated at 37°C for 15 minutes after filtering. Once the 15 minutes of incubation was complete, 50 µL of DNAse Solution (DNAse reconstituted from Worthington Kit with 500ug HBSS) was added to the organoids before gentle trituration using a wide bore tip. Mixture is incubated again for 10 minutes before being triturated with a p1000 tip and then a p200 tip. After this step, the solution was strained through a 70 µm cell strainer and then a 30 µm cell strainer. Another 1 mL of Neurobasal was used to rinse the cells prior to centrifugation at 300 rcf for 5 minutes. The supernatant that formed was removed, and the cells were resuspended in 1X PBS and 0.04% BSA. Cells were kept on ice while concentration and viability were assessed with a hemocytometer (Bio-Rad). The volume of cells was transferred to a 1.5 mL microcentrifuge tube for single-cell RNA sequencing (sc-RNA).

#### 10XscRNASeq Protocol

Following organoid dissociation and cell count, cell concentration was adjusted to 700-1200 cells/µL for 10X single cell sequencing. For each sample, 7,500 cells were loaded into the 10X Chromium controller to target recovery 3,500 cells and Gel Beads in Emulsion (GEM) was generated. 10X Genomics 3’v3.1 chemistry was used, and the samples were processed following the 10X Genomics protocol. Single cell libraries were sequenced using the illumine NovaSeq 6000.

#### Single cell data alignment

10x sc-RNA sequencing data in Fastq files were aligned to transcripts using Cell Ranger 3.1.0 (). Reference genome GRCh38 (Ensembl 93) was used as the reference genome. In the CellRanger count command, parameters chemistry and expected cells were set as SC3Pv3 and 6000, respectively.

#### Single cell data processing and normalization

Seurat 4 was used analyze raw data from Cell Ranger h5 files. We filtered out low-quality cells by removing cells with less than 1000 or more than 10000 unique genes. To further remove low-quality/dying cells we removed cells with more than 15% mitochondrial transcripts. After quality control, we harvested 26,329 genes and 3366 high-quality cells from MIBOs organoid. Original and processed data can be found in Data Availability. We applied the default normalization approach of Seurat.

#### Cell Annotations

Following quality control, we found a total of 22 clusters generated with a fine resolution (2.0) from dissociated MIBOs. Top 100 Differentially Expressed Genes (DEGs) for each pre-annotated cluster was found using ‘FindAllMarkers’ and can be found in Supplemental Table 1. We performed a manual annotation for each cluster based on canonical markers from precious studies, such as NES and HES1 for apical progenitors, ASCL1 and HES6 for basal progenitors, and STMN2, MAP2 and DCX for neurons. To assist in cluster annotation, we filtered ribosomal genes post clustering. We then used canonical telencephalic markers to combine initial clusters based on similarity in gene expression profile. A total of 10 cell classes were derived and included various subpopulations of GE derived cells. Cluster 15 was labeled as unknown given that there was no clear association based on expected marker genes.

#### Device Fabrication

Individual pieces for MIBOs device were made entirely out of PDMS as previously described by Li et al., 2021. Modifications included a PDMS Matrigel embedding area which used a PDMS film and laser cut (40 W Epilog Mini 18 x 12) three ascending positions at 20% speed, 80% power, 2500 Hz frequency, and 600 DPI resolution with vector job type (Fig1, Fig S1). Additionally, two thin PDMS slabs (5cm x 2cm) are glued to the front and back the device frame to form a self-contained area for culture medium.

### Quantification and Statistical Analysis

#### MATLAB Gradient Quantification

We used MATLAB and its inbuilt operators and functions to develop a program capable of analyzing the local change in pixel intensity. The program was then run on raw images captured by Leica DMi8 microscope. Here we leveraged the ‘Sobel’ gradient operator which uses the weighted sum of pixels in a 3x3 neighborhood to determine the gradient direction and gradient magnitude of the pixel. We also used the ‘quiver’ function to visualize the gradient direction (angle of quiver) and gradient magnitude (size of quiver) of individual pixels within an image. The sum of each pixel’s gradient magnitude was used for our analysis (Fig 2B, 2C). Image intensity was normalized by dividing all analyzed pixels by the value of the brightest pixel in their respective image providing a range of 0-1. An individual binary mask was made to assure that only regions of our organoid sections were measured and that any IHC artifacts were limited. MATLAB code available: https://github.com/Npavo002/MATLAB_SobelGradientAnalysis

#### Quantification and Statistical Analysis of Calcium Imaging Data

Data wrangling was done in Microsoft Excel with raw data points transferred to Prism (9.3.0) for basic statistics, outlier detection, significance tests, and graph generation. A ROUT outlier test was performed to identify outliers within each data set. The data was fitted with nonlinear regression and a false discovery rate of Q=1%. To test for statistical significance a one-way ANOVA test with multiple comparisons (Dunnett’s test) was used to compare conditions.

