## Supplemental information for "Patterning ganglionic eminences in developing human brain organoids using morphogen gradient inducing device"

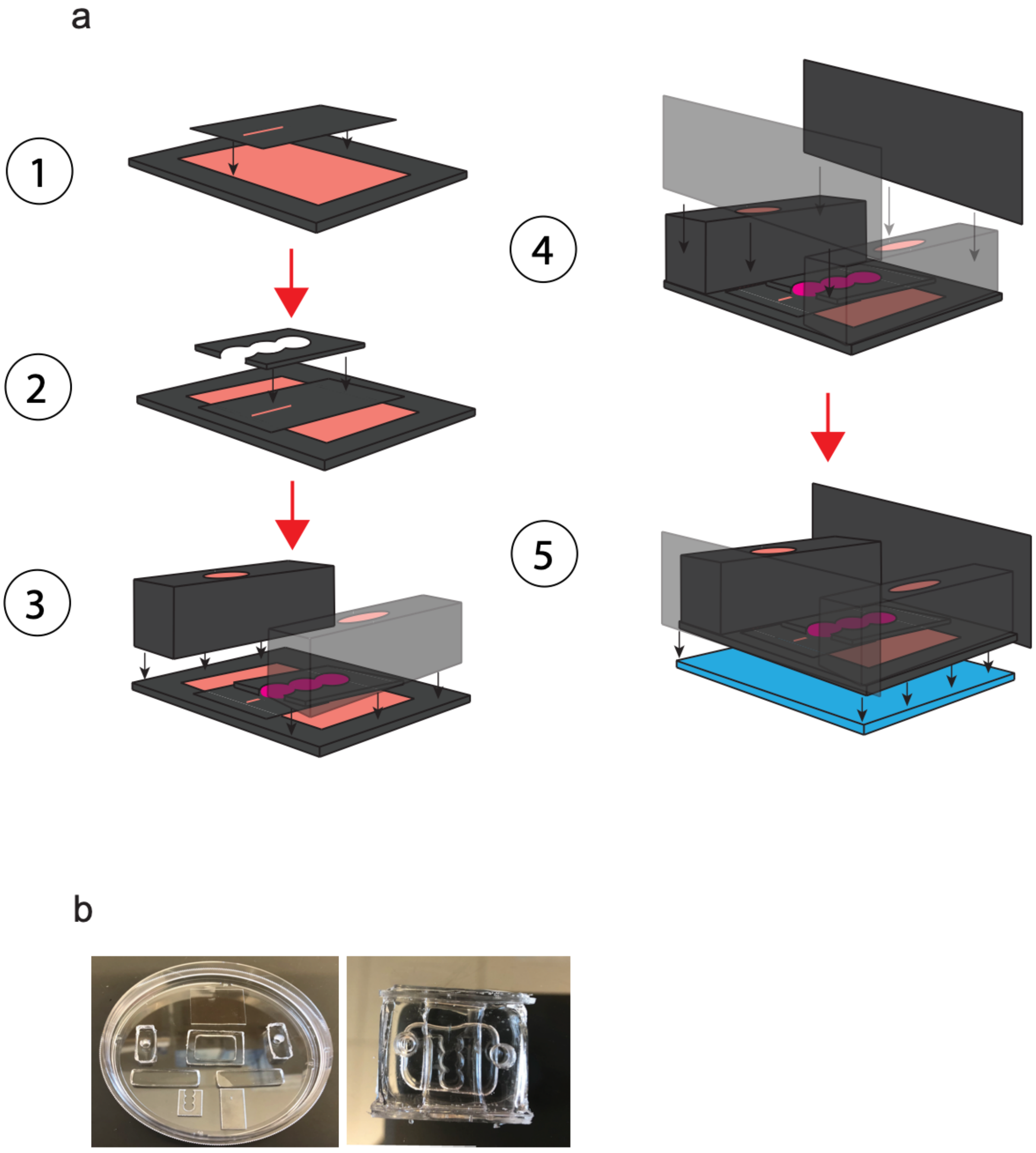


**Supplemental FIGURE 1 | PDMS device assembly.**

(A) Numbered, step by step schematic of PDMS device assembly. 1: Culturing area with laser-cut slit placed on chemical chamber base. 2: Matrigel embedding scaffolds placed over culturing area. 3: Inlet/outlet placed on both sides of device. 4: Encapsulating pieces to create a self-containing region for culture medium. 5. Entire device is secured to a glass bottom to fully seal the chemical reservoir.

(B) (Left) Unassembled, Individual PDMS slabs required to make one PDMS device. (Right) Fully assembled PDMS device.

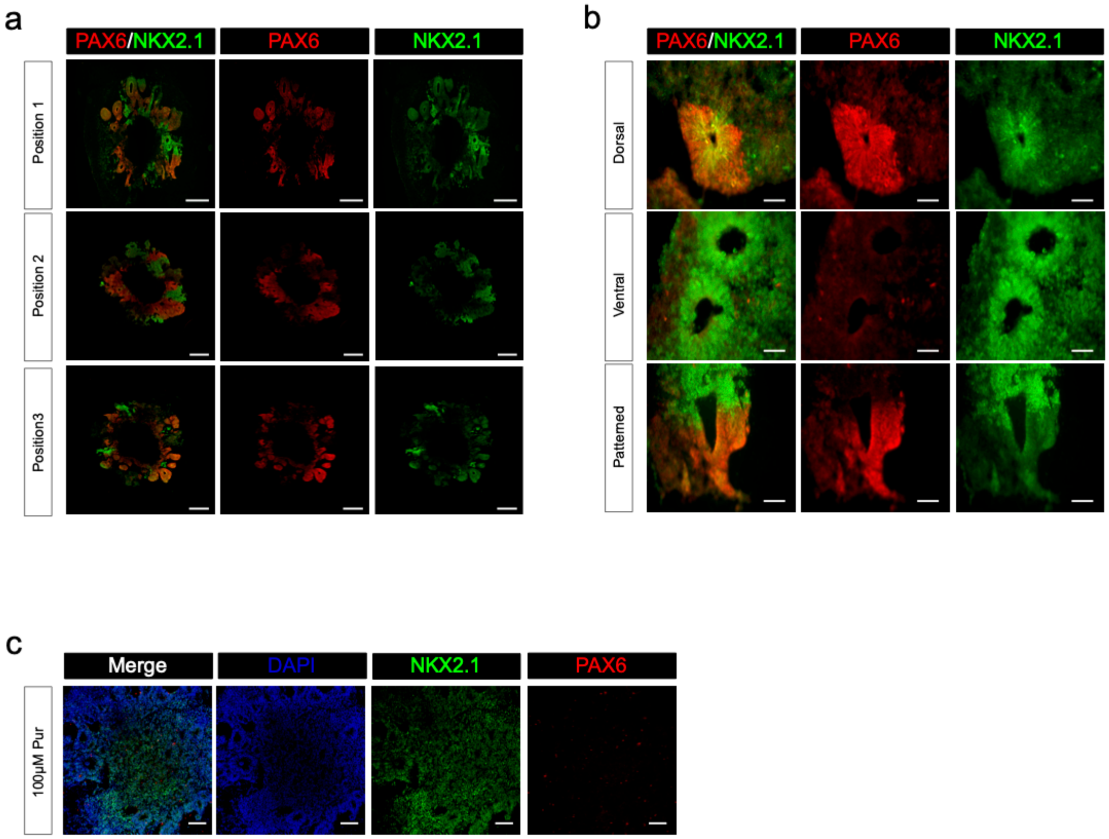


**Supplemental FIGURE 2 | Characterizing the effects of purmorphamine (Pur) gradient on forebrain organoids.**

(€) Representative confocal images of NKX2.1 and PAX6 staining in device-patterned organoids at different distances from Pur source. 500 µm scale bars.

(B) Representative confocal images displaying various rosette features observed in patterned organoids. 50 um scale bars.

(C) Organoid cultured in PDMS device and exposed to 100uM Pur gradient shows full ventralization. 200µm scale bar.

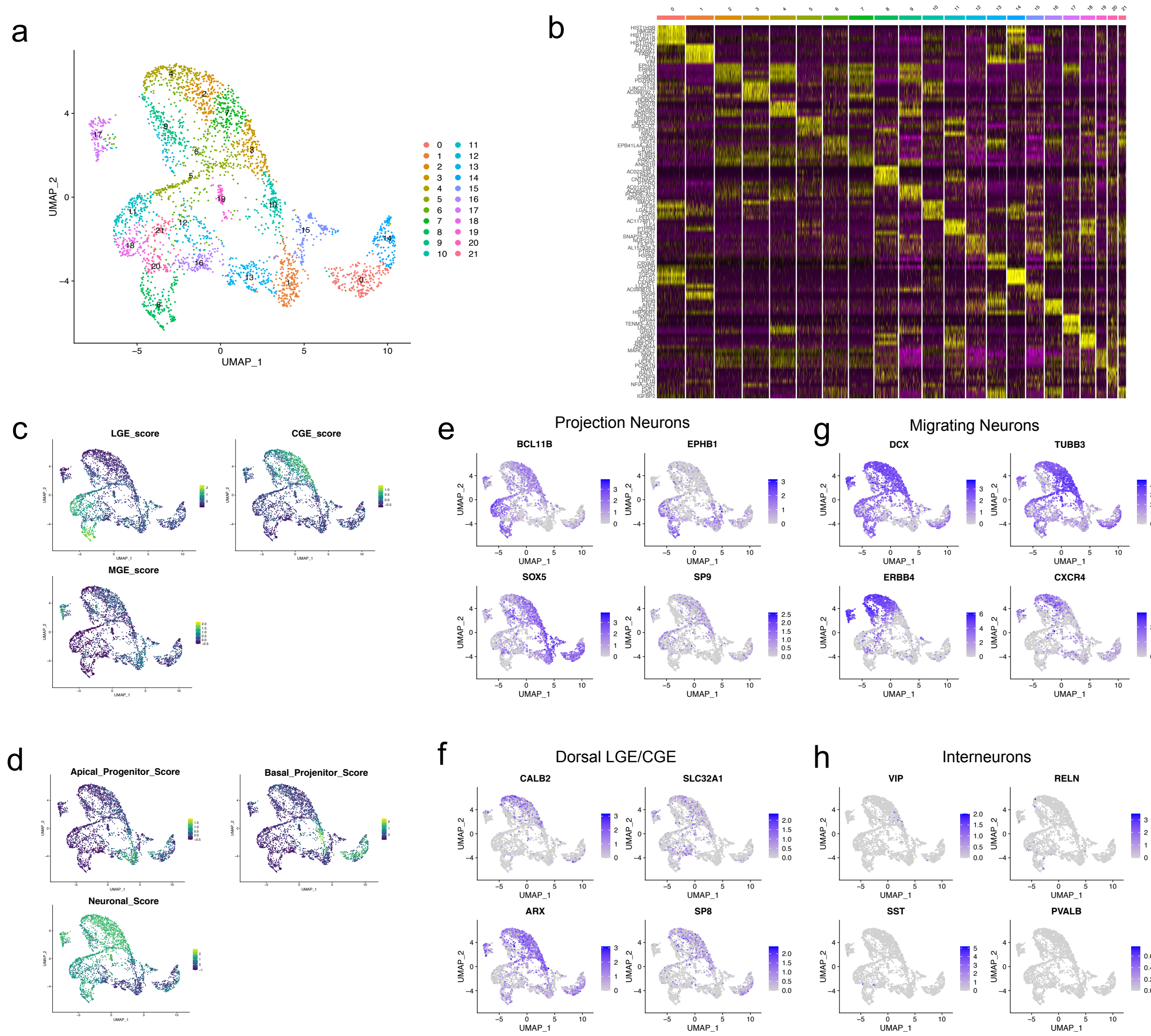

**Supplemental FIGURE 3 | Characterization of mature neuronal subtypes in 4-month device grown organoids.**

(A) UMAP of 3,366 single cells dissociated from 4-month MIBO before cell annotation. A total of 22 clusters were established using the first 30 principal components and a fine resolution of 2.0.

(B) Heatmap showing top 5 genes expressed for each cluster. €) UMAPs showing combined expression for canonical markers for each of the GEs. LGE: ZFHX3, ZNF503, MEIS2, FOXP1, EBF1, ISL1, RBFOX1. CGE: NR2F2, PROX1, SP9, SCGN, PRKCA. MGE: NKX2.1, LHX6, LHX8, SOX6.

(D) UMAPs showing combined expression for progenitors and mature cell types. Apical progenitors: NES, HES1. Basal progenitors: ASCL1, HES6. Neuronal cells: MAP2, STMN2, D€

(E) Feature maps for key marker of projection neurons.

(F) Feature maps identifying dorsal LGE and CGE.

(G) Feature maps showing migrating neurons with top row for general migration and bottom row more specific to IN migration.

(H) Feature map displaying presence detected for CGE and MGE derived INs.

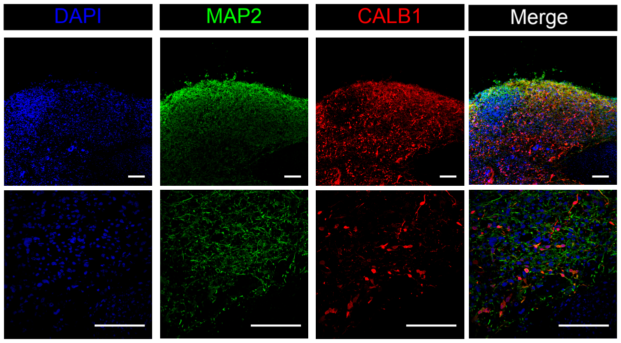


**Supplemental FIGURE 4 | Confirmation of CALB1+ interneurons.**

(A) Representative confocal images of mature MAP2+ neurons with CALB1+ migrating interneurons in in 3.5-month MIBO sections. Scale bar is 100um for both objectives.
